# Personality heterophily and friendship as drivers for successful cooperation

**DOI:** 10.1101/2023.09.19.558534

**Authors:** Debottam Bhattacharjee, Sophie Waasdorp, Esmee Middelburg, Elisabeth H.M. Sterck, Jorg J.M. Massen

## Abstract

Cooperation is widespread and arguably a pivotal evolutionary force in maintaining animal societies. Yet, proximately, what underlying motivators drive individuals to cooperate remains relatively unclear. Since ‘free-riders’ can exploit the benefits by cheating, selecting the right partner is paramount. Such decision rules need not be based on complex calculations and can be driven by cognitively less-demanding mechanisms, like social relationships (e.g., kinship, non-kin friendships, dyadic tolerance), social status (e.g., dominance hierarchies) and personalities (social and non-social traits); however, holistic evidence related to those mechanisms is scarce. Using the classical ‘loose-string paradigm’, we tested cooperative tendencies of a hierarchical primate, the long-tailed macaque (*Macaca fascicularis*). We studied three groups (*n=32*) in social settings, allowing free partner choice. We supplemented cooperation with observational and experimental data on social relationships, dominance hierarchies, and personality. Friendship and dissimilarity in a non-social ‘exploration’ personality trait predicted the likelihood of cooperative dyad formation. Furthermore, the magnitude of cooperative success was positively associated with friendship, low rank-distance, and dissimilarity in an ‘activity-sociability’ personality trait. Kinship did not affect cooperation. While some findings align with prior studies, the evidence of (non-social)personality heterophily promoting cooperation may deepen our understanding of the proximate mechanisms and, broadly, the evolution of cooperation.

## 1. Background

Cooperation, defined as the collaborative efforts of two or more individuals to achieve a common goal [1], is widespread in the animal social world and may exist in diverse forms, from breeding to hunting [2,3]. While humans are hyper-cooperative and exhibit incredibly complex dynamics of cooperation [4], several non-human animals also demonstrate surprisingly high intra- and interspecific cooperative tendencies [5]. Cooperation can serve as the building block for establishing and sustaining societies and may offer fitness benefits to group members [1,5,6]. Although the evolutionary mechanisms of cooperation are well studied from theoretical perspectives [7], what motivates individuals to cooperate with a partner remains poorly understood from a proximate mechanistic point of view. In other words, the question remains: How do individuals choose their cooperative partners? This is particularly important because cooperative interactions can present exploitable benefits. ‘Free-riders’ can reap the benefits without any contributions, thereby hampering the sustenance of cooperation [8]. While punishment rules can be enforced to stop exploitation [9,10], they may require considerable cognitive resources. Therefore, an efficient and preventive step would be to select reliable partners through ‘cognitively less demanding’ decision rules. Non-human animals may adopt various cooperative decision rules and employ suitable strategies to choose reliable partners. However, a comprehensive understanding of these mechanisms and their influence on partner choice remains limited due to the lack of holistic examination and the predominant use of ‘forced dyads’ in most experimental studies on cooperation.

Kin selection elucidates the basis for cooperative interactions among genetically related individuals [11]. Nevertheless, cooperation extends its reach beyond the confines of kinship and may endure through reciprocity among non-kin [12]. Members within a social group usually have ample opportunities for interactions, facilitating the occurrence of reciprocity. As a result, strategies such as ‘(generous) tit-for-tat’, ‘win-stay’, and ‘lose-shift’ may effectively operate within networks of repeated interactions to foster cooperation [13,14]. Conventionally, the cognitively sophisticated ‘calculated reciprocity’ was thought to drive cooperation [15], where cooperative investment is contingent on individuals’ mental bookkeeping of the costs and benefits of interactions. A lack of empirical support for this reciprocity in animals due to several cognitive constraints (see [16]) led researchers to propose simpler mechanisms, like ‘attitudinal-’ [17] or ‘emotional reciprocity’ [18]. These forms of reciprocities suggest that individuals may develop emotionally mediated attitudes toward group members through social interactions [19–22], which help them make cooperative decisions. In addition, the relatively simple ‘symmetry-based reciprocity’ suggests that individuals can engage in reciprocity based on symmetrical variables like age, dominance rank, kinship, etc. [15]. However, this hypothesis did not receive much support as reciprocity sustains even when the symmetrical variables are ruled out [21,23]. Although there is ambiguity regarding which of these mechanisms best predicts reciprocity [24], decision rules with low cognitive demands garnered the most support, which can further aid in detecting cheaters or avoiding potentially defecting partners, thereby increasing cooperative success in non-human animals.

Social tolerance, defined by an individual’s willingness to allow others in proximity while obtaining a high-quality reward [25], has significant implications for cooperation. Enhanced social tolerance can drive prosocial and cooperative tendencies [4,26,27]. Therefore, varying dyadic tolerance levels within a group can determine the formation and maintenance of cooperative interactions. Like tolerance, friendship is widely acknowledged as a catalyst for cooperation [28–30]. Although, in a stricter sense, strong non-kin affiliations are considered friendships, the most commonly found friendships are kin-based [31]. Yet, kinship may facilitate bond formation but may not be a prerequisite for friendship [32]. Moreover, social tolerance and friendships can work independently despite having similar underpinnings. For instance, it is far-fetched to say that socially tolerant individuals are always friends. But friendship is typically driven by enhanced tolerance. The positive effect of friendship on cooperation success aligns with the idea that strong social bonds can enhance the fitness of the partners [33,34]. While this preferential selection may create a ‘biological market’, where friends are associated with assured benefits [35], choosing a friend is also more predictable and cognitively less demanding than choosing a stranger or non-friend while collaborating [36]. Thus, differentiated strengths of dyadic tolerance and friendships can play a crucial role in forming cooperative dyads and in their maintenance and success in complex societies. Yet, empirical studies investigating the direct influence of friendship on cooperation in non-human animals are lacking.

Another key proximate mechanism of cooperation is consistent inter-individual behavioural differences or personalities [37]. They are fundamental behavioural constructs that distinguish one individual from another and hold considerable evolutionary significance [38,39]. Empirical evidence suggests that similarity in personality, or personality homophily, is a strong predictor of friendship [40–43], which in turn may foster cooperation. People with ‘prosocial personalities’ like agreeableness are more likely to cooperate [44]. Similarly, extraversion and cooperative success in humans are linked positively (see [45]). A few studies on non-human primates so far show similar patterns: Barbary macaques (*Macaca sylvanus*) with similarities in the shyness-boldness personality axis cooperate with each other [46], and personality similarities are predominant in cooperatively breeding species like common marmosets (*Callithrix jacchus*) [47]. Apart from primates, assortative mating based on personality similarity is linked to higher reproductive benefits in birds. For instance – a cross-fostering breeding experiment on zebra finches (*Taeniopygia guttata*) found that personality similarity within breeding pairs positively affects offspring fitness [48]. In cockatiels (*Nymphicus hollandicus*), higher as compared to lower behavioural compatibility between mates is linked to better-coordinated incubation [49]. Interestingly, personality traits positively affecting cooperation, such as extraversion, sociability, and agreeableness, are often grouped as ‘cooperative personality’ in humans [50]. In contrast to personality homophily, humans with dissimilar personalities can effectively engage in successful cooperative interactions, too. Cooperating individuals, each with a specific personality, may bring different problem-solving perspectives [51], thereby enhancing group performance [52,53]. Again, there have been reports about similar patterns in non-human animals, albeit scarcely. For example, rooks (*Corvus frugilegus*) successfully cooperate when dyads consist of a shy and a bold individual rather than two shy individuals [54]. In chimpanzees (*Pan troglodytes*), ‘impact’ males, dissimilar to the rest of the group, work as hunting catalysts [55]. Furthermore, disassortative mating strategies can be beneficial; they are often based on personality differences. Female rainbow Kribs (*Pelvicachromis pulcher*) choose males of dissimilar levels of boldness [56], and great tits (*Parus major*) with varying explorative tendencies are more likely to be in a partnership [57]. While the prevailing hypothesis revolves around personality homophily as a predictor of cooperative success, the potential impact of personality dissimilarities on cooperation remains largely overlooked and consequently unexplored, especially in non-human animals. Nevertheless, it appears that while homophily in social personality traits (e.g., agreeableness, sociability, affiliation, cf.[58]) is predictive of cooperative success, heterophily in non-social traits (e.g., exploration, boldness, activity) may facilitate cooperation. However, this remains a testable hypothesis to address whether and how individuals with personality (dis)similarity actively select one another as cooperative partners. Consequently, our understanding of how personalities influence cooperative decision-making and partner choice remains obscure.

Here we investigate the cooperative tendencies of a group-living primate species that has a matrilineal hierarchical social structure, the long-tailed macaque (*Macaca fascicularis*). While the existing covariation framework might suggest that a steep hierarchical structure could hinder cooperation [59], a growing number of empirical studies provide evidence of prosocial and cooperative tendencies in despotic societies [60–66]. Prosocial and cooperative behaviours seem to be selectively maintained through interdependence in these hierarchical systems [66]. Besides, rank similarities, i.e., low rank distances, can also be a strong mechanism favouring cooperation when hierarchies are relatively steep (see [67]). We employed a ‘loose-string paradigm’, wherein two individuals can attain rewards only by simultaneously pulling two loose ends of a single string [68]. Importantly, our study took place within the macaques’ existing social groups, thereby allowing for free partner choice. While we did not explicitly assess partner choice by manipulation, any discernible pattern from our free partner choice set-up is expected to indicate the underlying mechanisms efficiently. In addition to cooperation, we experimentally measured the long-tailed macaques’ dyadic tolerance levels. Finally, we supplemented cooperation and tolerance with data on friendships, dominance hierarchy and personalities, collected using extensive behavioural observations and experimental assays.

We hypothesise that individuals will use cooperative decision rules based on kinship, social tolerance, friendships (marked by contact sit, including both kin- and non-kin-based friendships), and dominance rank relationships. We predict that the likelihood of cooperation and its success would be positively associated with kinship, social tolerance, and friendship, whereas a negative effect of disparity in dominance ranks is expected. Finally, we hypothesise that personality will play a significant role in shaping both the establishment and success of cooperative dyads. Nonetheless, due to complex personality trait characteristics, we put forward a two-tailed prediction; we expect personality homophily and heterophily to influence cooperation at the trait level independently. In particular, we predict that homophily in affiliation and sociability-like traits (i.e., social personality traits) and heterophily in exploration and activity-like (i.e., non-social personality traits) traits will be positively linked to cooperation and its success.

## 2. Methods

### 2.1 Subjects and housing

We studied three social groups of captive long-tailed macaques residing at the Biomedical Primate Research Centre (BPRC) in Rijswijk, the Netherlands. The first group (Gr.1) consisted of 15 individuals, out of which four were under the age of one year at the beginning of the study and thus not included (n = 11). Among the remaining 11 individuals were seven adult females and one adult male, all above three years of age, as well as two juvenile females and one juvenile male between one and three years old. The second group (Gr.2) originally had 18 individuals but was reduced to 17 as one male was removed early during the study due to compatibility issues with conspecifics (n = 17). The remaining 17 individuals comprised ten adult females, one adult male, two juvenile females and four juvenile males. The third group (Gr.3) consisted of four adult males (n = 4). The study was conducted in the macaques’ existing social groups and home enclosures. We neither separated the individuals from their social groups nor made forced dyads. The description of the individuals is provided in **table S1**, electronic supplementary material.

All groups had an indoor and an outdoor enclosure, and individuals could freely move within and between them. The enclosures of Gr.3 (indoor = 3.55 m^2^, outdoor = 3.88 m^2^) were considerably smaller in area compared to the enclosures of Gr.1 and Gr.2 (indoor = 49 m^2^, outdoor = 183 m^2^). The study groups had no visual contact with each other as they were kept in different buildings or on different floors of the same building. All enclosures had multiple enrichment structures, including slides made from firehoses, plastic toys, wooden structures of various heights, platforms, climbing stairs, a plastic pool (except for Gr.3), and a tree trunk. Extra enrichment materials, such as branches and paper containers, were provided when available. The indoor enclosure had concrete floors covered with sawdust bedding, while the outdoor enclosures were covered with natural soil and sand substrates. The feeding of the animals always took place in the indoor enclosures. The diet consisted of monkey pellets in the morning, placed in feeding cans attached to the enclosure fences, and vegetables in the afternoon, with an occasional seed mix (corn, sunflower, etc.) being thrown in to stimulate foraging behaviour. Water was available 24/7 ad libitum in both indoor and outdoor enclosures. Animals were never food- or water-deprived for our experiments, and the normal feeding schedule was followed on each testing day.

### 2.2 Experimental design and data collection

#### 2.2.1 Loose-string paradigm

We conducted a ‘loose-string paradigm’ to assess cooperation in this study [69,70]. The experimental apparatus consisted of a sliding platform (width = 1.1 m) on top of a larger wooden base (width = 1.3 m). The sliding platform had two feeding trays made of plastic positioned on each end of the section that faced the enclosures. The placement was done in such a way that prevented one individual from reaching both feeding trays simultaneously. The opposite section had a hard plastic handle attached, which the experimenter could pull to move the platform. On the side of the handle, two metal loops were anchored. Depending on the experimental phase, the string(s) were either attached to or inserted through the metal loops (see below). Throughout the experiment, corn, sunflower seeds or peas were used as food rewards alternately, as advised by the in-house veterinarians. The macaques were highly motivated to receive each of these items as a reward.

In Gr.1, the experiment commenced between November 2021 and early February 2022. We tested Gr.2 between July and October 2021. Gr. 3 was tested between June and August 2021. Experiments were conducted outdoors for Gr.2 and Gr.3, whereas, due to low temperatures during the winter months, the experimental setup was placed indoors for Gr.1. We tested the macaques one to three days a week in their existing social group settings, and experiments lasted a maximum of three hours per day. The cooperation experiment consisted of three different phases (c.f. [69,71]) – (i) habituation, (ii) training and social tolerance, and (iii) testing. The participating individuals were identified for all trials in the following phases.

i. *Habituation* – This phase allowed individuals to familiarise themselves with the apparatus and eliminate any potential bias of neophobia. The apparatus was placed in front of the enclosure, and mixed food rewards were placed on the feeding trays. The experimenter gently moved the platform towards the macaques and waited for them to obtain the rewards. Individuals who obtained the rewards at least three times were considered habituated. Furthermore, the following criterion was followed: at least half of the individuals from a group needed to be habituated to proceed to the next phase. The habituation phase lasted for two consecutive days.
ii. *Training and social tolerance* – Two separate strings were attached to the metal loops in this phase. Thus, pulling either string by one individual could move the platform close for obtaining food from the feeding trays. On the one hand, this phase allowed individuals to self-train themselves with the pulling mechanism, and simultaneously, we also measured the tolerance of the macaques to their group members. A trial began when both feeding trays were baited with a food reward (always the same type of food). The experimenter called “monkeys” to get attention and presented the strings to them. We recorded how often individuals retrieved food rewards near their group mates at the other string. A trial ended when both rewards were obtained or after 2 minutes. If the individuals did not pull for three consecutive trials, the session was called off and resumed the next day. We conducted a total of 18 sessions per group, with each session having 20 trials. A maximum of four sessions were carried out a day. The inter-trial and inter-session intervals were 20 seconds and 5 minutes, respectively. After completing all sessions, we checked if at least half of the individuals from a group pulled and obtained rewards in a minimum of 10 trials to proceed to the next phase. All three groups met these requirements, so all three could proceed to testing.
iii. *Testing* – We tested the cooperative behaviour of the macaques in this phase. Instead of two strings, one string was inserted through the metal loops, and the loose ends were presented. Therefore, the individuals had to simultaneously pull the two loose ends to move the platform and access the food rewards (electronic supplementary material, **movie S1**). If one individual pulled, the string became unthreaded, and the platform did not move from its initial position. A total of 30 sessions were conducted per group, each with 20 trials. Again, a maximum of four sessions were carried out a day, and inter-trial and inter-session intervals were 20 seconds and 5 minutes, respectively.

D.B. tested Gr.1, and S.W. performed the tests at Gr.2 and Gr.3. The macaques were familiar to both experimenters. We recorded the study using Canon Legria HF G25 and Sony FDR AX100E 4k video cameras mounted on tripods. Moreover, for Gr.2 and Gr.3, an instant coding sheet was maintained and later compared with the videos.

#### 2.2.2 Personality

The personality scores of the individuals were the same as in a previous study [72]. Here, we briefly summarise the methods and findings for convenience. We took a multi-method approach of behavioural observation and experiments to assess long-tailed macaques’ personality traits (also see [73]). A ‘bottom-up’ approach was used to avoid any subjective bias of predetermined clustering of behavioural variables. All 32 individuals tested in the current study were observed in two phases (with at least two months of interval) using 20-min long continuous focal follows (Phase 1: mean ± standard deviation = 159.76 ± 2.67 observation minutes/individual; Phase 2 = 159.89 ± 2.64 min/individual) in their existing social group settings. In addition, novelty experiments were conducted during the same phases to capture rare behaviours, like task solving, persistence, and exploration. Food puzzles (pipe and wooden maze), novel food items (dragon fruit and rambutan/starfruit) and novel objects (egg container and massage roller) were repeatedly used, again with at least two months of intervals in between. As personality traits are considered inter-individual differences that are consistent over time [37], the repeatability of the variables was checked between the two phases using an intraclass correlation analysis (ICC(3,1)). A subsequent principal component analysis provided us with three non-correlating dimensions or personality traits consisting of highly repeatable variables. We named the three dimensions ‘Activity-sociability’, ‘Affiliation’, and ‘Exploration’. Activity-sociability consisted of the following variables – *approach, leave, follow, pass by, leave passive, sit*, and *social play*; except for *sit*, all other variables were loaded positively. Affiliation had two positively loaded variables: *proximity* and *groom*. Finally, the personality trait exploration included positively loaded *object manipulation, handling container*, and *hang* variables. Affiliation and exploration implied social- and non-social traits, respectively. It is worth noting that we labelled the trait activity-sociability to incorporate activity- and sociability-related variables. With the exception of *social play*, all other variables could account for the activity component without necessitating direct social interactions. Consequently, this particular trait, even though labelled as activity-sociability, to a large extent, can be attributed to the general activity of the individuals. Individual personality scores from each trait were extracted.

#### 2.2.3 Dominance hierarchy and friendship

Dominance hierarchies were known and reported in [72] for all three groups. A Bayesian Elo-rating method [74] was used to construct the hierarchies based on unprovoked submissive behaviours - *avoid, be displaced, fear grimace, flee*, and *social presence*, recorded during the focal observations described above. These variables were independent of the variables used for personality assessment. Individuals were placed according to their ordinal ranks in the hierarchy for the three groups separately. In addition, we used *contact sit* as a proxy for friendships (cf. [43]). It was defined as two individuals sitting or lying together with body parts other than limbs physically touching. This durational variable was also independent of the personality trait-related variables. From continuous focal observations (see [72]), the duration of contact sit was extracted for all within group dyads.

### 2.3 Data preparation

#### 2.3.1 Cooperation

Cooperation (also synonymously used with cooperative success) was defined as when two individuals in the testing phase pulled the two loose ends of the string and obtained rewards without monopolisation. It was coded as a binary variable (yes/no). Furthermore, we counted the number of times individuals in a particular dyad cooperated. Two experimenters (D.B. and S.W.) coded the cooperation data. Based on 12% of the data, inter-rater reliability was calculated and found to be excellent (Cohen’s kappa = 0.98). As mentioned, after the cooperation studies, a three-year-old male had to be separated from Gr.2 and thus was not assessed for personalities. We did not include the concerned individual in the cooperation data analyses. Consequently, all trials were removed where this individual was present. Unlike the 600 trials in the testing phase of the other two groups, we obtained only 459 trials from Gr.2. Similarly, the number of social tolerance trials differed between Gr.2 (192 trials) and the other two groups (each 360 trials). In Gr.1, Gr.2, and Gr.3, a total of 7 (participation rate = 63.6%), 10 (59%), and 4 (100%) individuals participated in the testing phase (i.e., at least in one trial), respectively. Theoretically, each participating individual had the opportunity to form a dyad with the other participating group members. In line with this, all potential combinations of dyads were constructed separately for the three groups (potential number of dyads: Gr.1 = 21, Gr.2 = 45, and Gr.3 = 6), and data on the likelihood and magnitude of cooperation were used for further analyses. Due to the zero-inflated nature of the data, a hurdle approach was considered to analyse cooperation, where we first looked at the likelihood (i.e., did a dyad cooperate at least once or not) and then the magnitude of cooperation (i.e., no. of successful trials when a dyad cooperated at least once; success > 0).

#### 2.3.2 Social tolerance

We used data from the social tolerance test phase to measure dyadic tolerance levels. Since we conducted the experiment in social group settings and individuals can have varying tolerance levels to their group members, our method considered all opportunities when two specific individuals could co-feed or monopolise. We counted the number of times two specific participating individuals obtained food in the presence of each other without any monopolisation (i.e., ‘co-feeding’ events). This was divided by the sum of those two individuals obtaining food alone (or monopolisation). The resulting values were normalised for each group and used as social tolerance scores. The tolerance score ranged from -1.08 to 6.05 (median = -0.21), with higher values indicating higher tolerance within a dyad.

#### 2.3.3 Personality differences

The absolute differences in personality scores were calculated separately for each dyad for the three traits. Activity-sociability, affiliation, and exploration had a range of 0.03 – 3.42 (median = 1.13), 0.02 – 4.86 (median = 1.01), and 0.008 – 3.05 (median = 1.13), respectively, with higher values suggesting greater dyadic personality differences.

#### 2.3.4 Friendship

We extracted the observed duration of contact sit for all within group dyads. The duration of contact sit per dyad was divided by the total observation time of a dyad to obtain the rate. Dyadic contact sit rates were further divided by the mean rate of a group [33,75]. Finally, the resulting values were z-transformed at the group level to get contact sit scores. A score range of -0.52 to 2.85 (median = -0.42) was obtained, with high scores indicating prolonged sitting in body contact, i.e., stronger friendships between dyad members than the average dyad in their corresponding group.

### 2.3 Statistics

All statistical analyses were conducted in R (version – 4.3.0) [76]. We first compared the number of trials in the testing phase where an individual pulled the rope in the presence versus absence of participating partners using a Wilcoxon Signed Rank test. This step ensured whether the participating animals understood the mechanism, i.e., the requirement of a partner to obtain rewards successfully. The proportion of successful cooperation across sessions was checked for the three groups using Spearman rank correlations. Since personality homophily can predict friendships, we first checked if there were significant correlations of contact sit scores with each personality trait using Spearman rank correlations. Similarly, we performed a correlation test between contact sit scores and dyadic social tolerance. Generalised linear mixed-effect model (GLMM) analyses were performed to investigate the likelihood and magnitude of cooperation. The first model (for likelihood) had cooperation success as the response variable (yes/no) with a binomial error distribution. In contrast, the other model had the number of such successful cooperation trials (i.e., magnitude, success > 0) as the response variable and used a Poisson error distribution. Additionally, this model included the number of trials in the testing phase as an offset term to account for the variations. Both models included the following fixed effects – dyadic personality difference scores (the three traits were included separately), dyadic social tolerance scores, friendship scores (i.e., dyadic contact sit scores), sex composition (female-female/female-male/male-male), age difference (in years), dominance rank distance, and kinship (yes/no; mother-offspring, siblings), and random effects – identities of the two individuals and identities of the dyads nested within groups.

GLMM analyses were conducted using the *glmmTMB* package [77]. Collinearity among the fixed effects was checked using the *performance* package [78], and a variation inflation factor (VIF) value of < 4 was considered as low or no collinearity. The null models lacked all fixed effects but retained random effects. Null vs. Full model comparisons were checked with the help of ‘lrtest’ function of the *lmtest* package [79]. We selected the best-fitted models based on Akaike Information Criteria (ΔAIC = 2 as threshold) values, and only the best-fitted models have been presented in the results. Model diagnostics (residual normality, dispersion, and outliers) were investigated using the *DHARMa* package [80]. The significance value (α) was set at 0.05.

## 3. Results

The percentages of successful cooperation were 45.5%, 44.6%, and 32.16% for Gr.1, Gr.2, and Gr.3, respectively. The two mixed-sex groups (Gr.1 and Gr.2) exhibited cooperation at a comparable level (Fisher’s Exact Test: p = 0.80) but higher than the smaller all-male (Gr.3) group (Gr.1 and Gr.3: p < 0.001; Gr.2 and Gr.3: p < 0.001). Overall, individuals pulled the string significantly more often with a partner than when alone (Wilcoxon Signed Rank test: z = 2.279, p = 0.02), suggesting that they seem to have at least some understanding of the need for a partner to obtain rewards. For all three groups, a positive trend or a significant correlation was found between the proportion of successful cooperation and test sessions (Spearman Correlation Test, Gr.1: rho = 0.30, p = 0.10; Gr.2: rho = 0.34, p = 0.06, Gr.3: rho = 0.79, p < 0.001), further indicating an effect of learning.

We found no evidence of a correlation between friendship and personality differences (activity-sociability: rho = -0.20, p = 0.09; affiliation: rho = -0.09, p = 0.44; exploration: rho = - 0.06, p = 0.59). Similarly, no correlation was found between friendship and dyadic social tolerance (rho = 0.002, p = 0.98). Therefore, instead of covariates, individual effects of personality, friendship, and tolerance were tested in the subsequent statistical models. Nonetheless, we checked for collinearity among these variables for further confirmation. Additionally, no difference in friendship between kin and non-kin members was found (Mann Whitney U Test: z = 1.011, p = 0.312).

Based on the best-fitted model, friendship and exploration trait differences predicted the likelihood of cooperation success. A positive association was found between the likelihood of cooperation success and friendship (GLMM: z value = 3.043, p = 0.002, **fig 1**). In addition, higher differences in exploration trait scores predicted the likelihood of cooperation (GLMM: z value = 2.914, p = 0.003, **fig 2**). No effect of activity-sociability, affiliation trait differences, social tolerance, and kinship were found. The best-fitted model differed significantly from the null model (Likelihood ratio test: χ^2^ = 25.58, p < 0.001). We found no collinearity among the fixed effects (VIF range = 1.42 – 3.53).

**Fig. 1.**
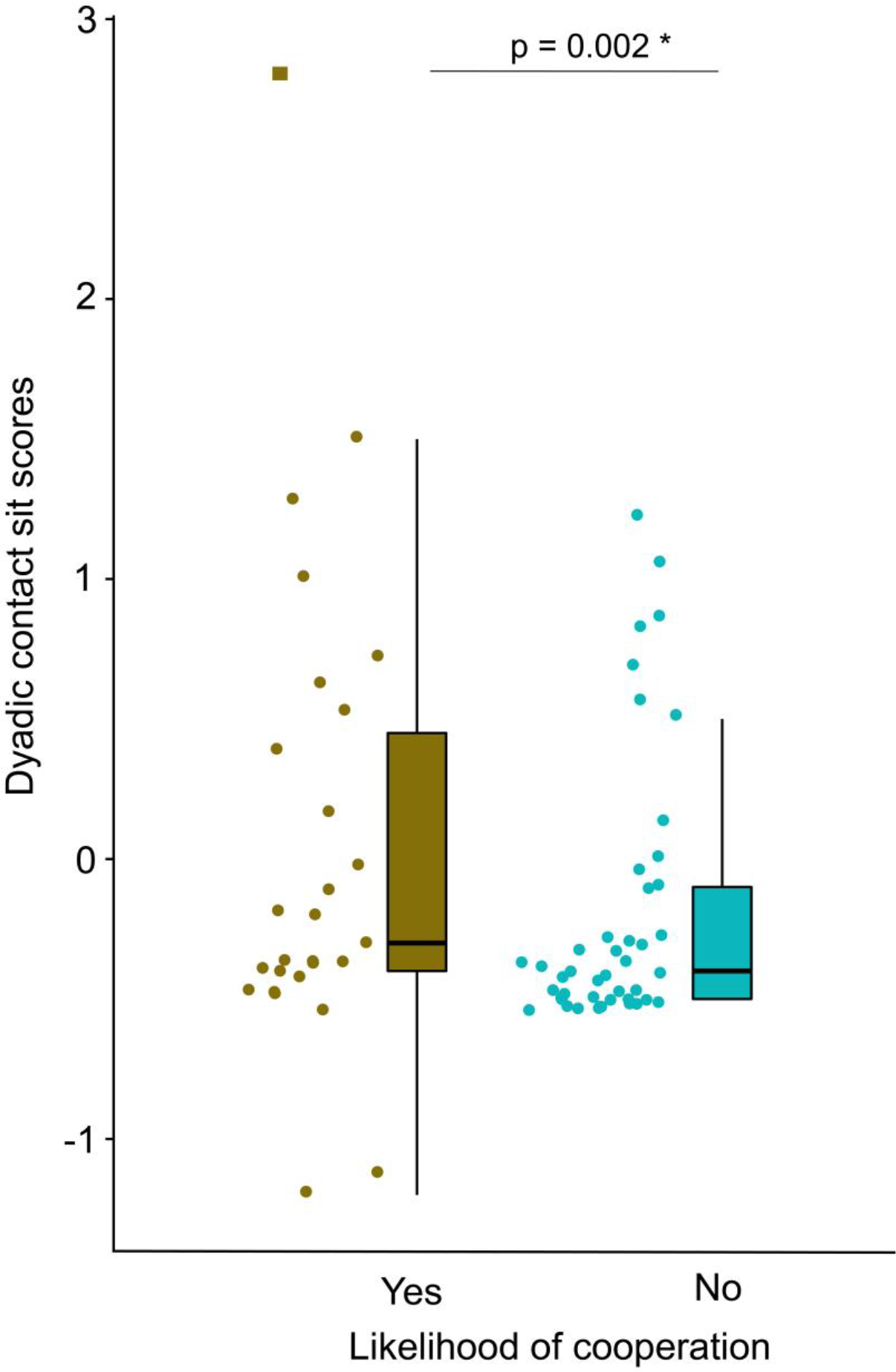
Dyadic contact sit and likelihood of cooperation. A box-whisker plot showing the contact sit scores for cooperative and non-cooperative dyads (GLMM: p = 0.002). Boxes represent interquartile ranges, and whiskers represent the upper and lower limits of the data. The horizontal bars within the boxes represent the median values. Outlier highlighted by square.

**Fig. 2.**
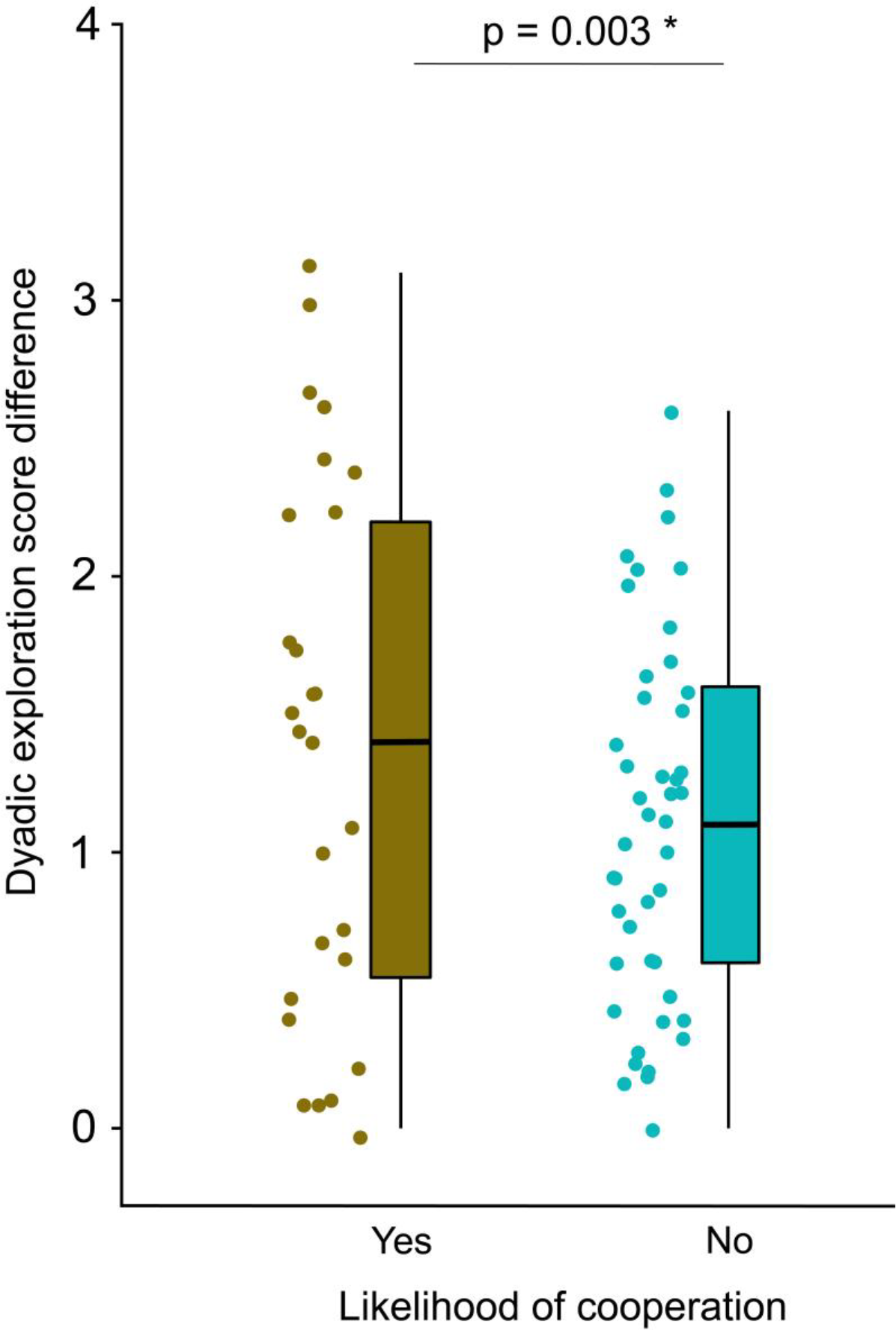
Exploration trait differences and likelihood of cooperation. A box-whisker plot showing the personality trait exploration score differences for cooperative and non-cooperative dyads (GLMM: p = 0.003). Boxes represent interquartile ranges, and whiskers represent the upper and lower limits of the data. The horizontal bars within the boxes represent the median values.

We found that friendship, dominance rank distance, and activity-sociability trait differences significantly predicted the magnitude of cooperation. A higher cooperation success was associated with friendship (GLMM: z value = 2.550, p = 0.01, **fig 3a**) and lower dominance rank distance (GLMM: z value = -2.571, p = 0.01, **fig 3b**). And, a larger difference in activity-sociability trait scores were linked to higher cooperation success (GLMM: z value = 4.150, p < 0.001, **fig 3c**). We did not find any effects of affiliation and exploration score differences, social tolerance, and kinship on the magnitude of cooperation. Age difference and sex composition fixed effects were dropped from the best-fitted model due to an initial high VIF value and model convergence issue. The best-fitted model differed significantly from the null model (Likelihood ratio test: χ^2^ = 23.02, p = 0.001). No collinearity among fixed effects was found (VIF range = 1.23 – 1.96).

**Fig. 3.**
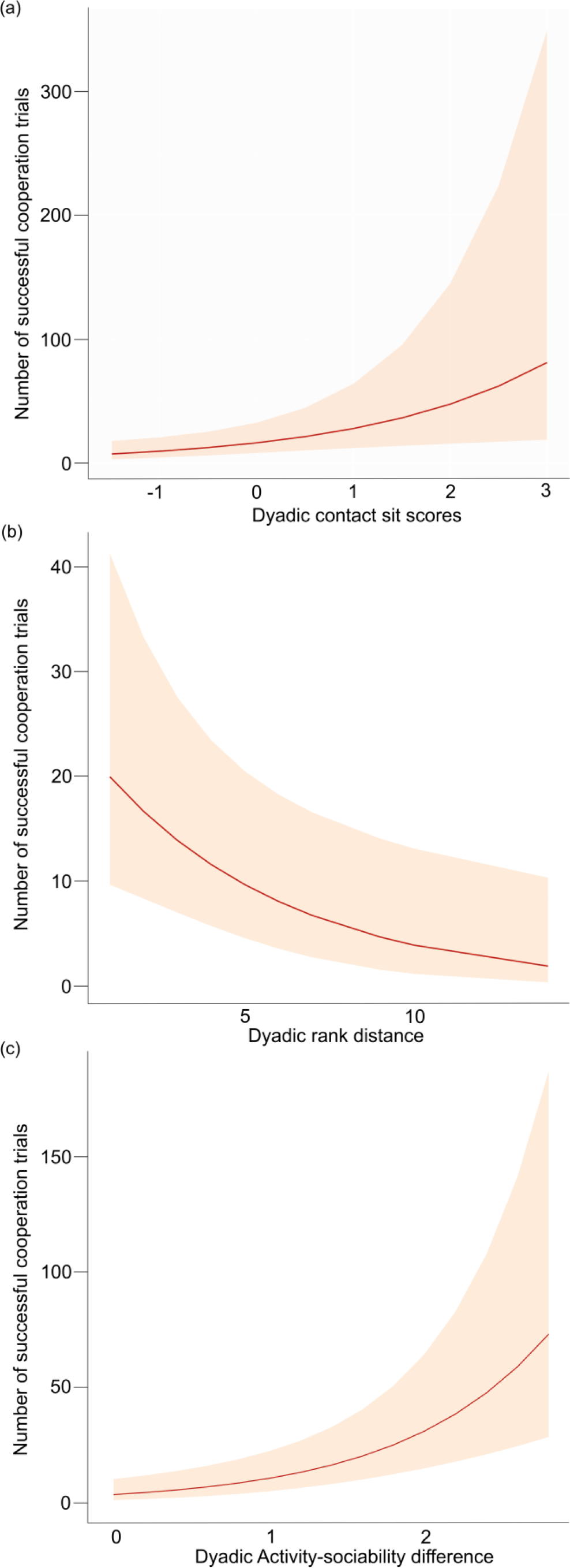
Effects of contact sit, rank distance, and activity-sociability differences on successful cooperation. (a) GLMM effect plot showing a predicted effect of dyadic contact sit on the number of successful cooperation, (b) GLMM effect plot showing a predicted effect dyadic rank distance on the number of successful cooperation, (c) GLMM effect plot showing a predicted effect activity-sociability personality difference on the number of successful cooperation. Shaded areas in the plots indicate 95% confidence intervals.

## 4. Discussion

We investigated some key proximate mechanisms of cooperation in a hierarchical group-living primate using an experimental setup that allowed free partner choice. Larger mixed-sex groups exhibited higher cooperation than a relatively smaller all-male group. Friendship predicted the formation of cooperative dyads and their success. Incongruence in dominance position had an inverse association with cooperative task success, suggesting that individuals with similar ranks in the hierarchy were more efficiently involved in cooperation. To our surprise, kinship and social tolerance did not influence cooperation. While we found no support for homophily in social personality traits fostering cooperation, heterophily in the non-social personality traits was linked to the formation of cooperative dyads and their success. This study provides valuable insights into the proximate mechanisms underlying partner choice in cooperation, allowing us to comprehend the evolution of cooperative interactions better.

Group-level variations in altruistic, i.e., prosocial and cooperative tendencies, are often ignored. Even though species-typical patterns are of utmost importance for understanding the evolutionary trajectories of cooperation, socio-ecological and ‘group-specific’ factors can contribute to considerable inter-group differences within a species (see [81]). The social compositions of our study groups, albeit in relatively smaller group sizes, represented long-tailed macaque social organisations typically found in the wild [82,83]; mixed-sex groups that include philopatric females from different matrilines represent a more stable form of social organisation than the bachelor male (i.e., dispersing sex) groups. This stability in social organisation might have resulted in higher cooperation in mixed-sex than in all-male groups. Nonetheless, even within all-male groups, strong social bonds may form (e.g., political coalitions), which may or may not be facilitated by kinship [32,84]. In general, hierarchical matrilineal societies exhibit kin bias, and thus, genetically related individuals may engage in nepotistic cooperation [85]. However, we did not find any evidence of nepotistic cooperation in this study. A previous study on long-tailed macaques found a positive association between kinship and prosocial tendencies [60]. Interestingly, when prosocial choices became costly, such associations seemed to disappear [64]. Constraints on nepotism, like the immediate exertion of competition within matrilines and the subsequent need for non-kin allies, can explain the lack of evidence of kinship influencing cooperation. In Japanese macaques (*Macaca fuscata*), non-kin alliances are formed and become more advantageous than kin-based allies, comprised of matriline members [86]. Similarly, the majority of the highly cooperative dyads are unrelated or distantly related in chimpanzees [87]. Finally, in a food-sharing experiment, long-tailed macaques do not discriminate between kin and non-kin members [88]. Hence, growing empirical evidence shows higher cooperative success among non-kin than kin members. But why non-kin dyads are more effective than kin dyads, and particularly in what settings, remain to be investigated. Nonetheless, taken together, kinship might not be the best predictor of cooperation in hierarchical primates; instead, its simpler underlying mechanism, like familiarity, may be at play when choosing a partner.

Incongruence in dominance rank can be detrimental to cooperation [89,90], and our study found support for this effect. The intensity of cooperation increased with decreasing rank distance. Non-human primates are known to assess their own and other group members’ social ranks [91]. Besides, in despotic societies, a relatively steep and linear dominance hierarchy provides clear predictability of the directions of agonistic interactions [92]. Therefore, cooperating more with individuals from similar dominance ranks can be a sustainable strategy to minimise exploitation and avoid potential conflicts (see [69]). However, the likelihood of cooperation in our study did not depend on rank distance, indicating an influence of other proximate mechanisms triggering cooperative dyad formation. This also suggests that repetitive interactions were necessary to determine with whom to cooperate more.

Our findings align with previous studies that friendship paves the path for cooperation [30,93,94]. Individuals were more likely to choose friends over non-friends to build cooperative dyads and subsequently became highly successful. However, a previous study on captive long-tailed macaques found no effect of social bonds on cooperative success [95]. Such a contrasting finding could be attributed to methodological differences and the use of forced dyads. Also, unlike contact sit in the current investigation, the previous study used proximity (and grooming instances) to measure social bond strengths. This potentially indicates that sitting in body contact is a more powerful proxy for friendship in non-human primates, especially in hierarchical societies, where grooming can be directed towards dominant individuals [43]. Moreover, we found that friendships were independent of kin relationships. Thus, choosing a group member as a cooperative partner seems to be driven by a simple decision rule of who sits next to whom but in body contact. From an evolutionary standpoint, on the one hand, cooperating with a friend may provide high returns by decreasing the risks of exploitation and cheating; on the other hand, it can enhance fitness benefits [34–36,93]. Even though the formation and maintenance of friendships may incur certain costs, the corresponding benefits are expected to outweigh them, resulting in an overall advantageous outcome for friendship to be selected as a viable strategy [93,94]. Interestingly, no substantial impact of social tolerance favouring cooperation was found. There are two possibilities: First, since social tolerance is a prerequisite for forming and maintaining friendships [43,71], an underlying enhanced tolerance among friends might have obscured the effects of our co-feeding measure. Second is the current methodology’s limitation in assessing tolerance among dyads that never were together at the training apparatus during the co-feeding assay. Empirically, we incline ourselves towards the latter as no correlation was found between dyadic social tolerance and friendships. However, it is essential to mention that the criterion for the social tolerance phase was satisfied. Nonetheless, we recognise the need for a more robust co-feeding tolerance measure in future investigations. For instance, adopting a ‘co-feeding plot’ approach could provide comprehensive information at both group as well as dyad levels [96].

Homophily in social personality traits, in this case, affiliation, exhibited no discernible impact on cooperation. Since we did not find any collinearity of affiliation with social tolerance or friendship, it might imply that the observed affiliation trait scores were contingent on partner-specific affiliative interactions. It would be worthwhile to investigate whether this selective yet discriminative mechanism is exclusive to despotic societies due to interdependencies (see [66]). Therefore, in socially tolerant species, homophily in social personality traits, in general, can be expected to act as a stronger and more efficient force driving cooperation. An alternate explanation would be the strong effect of friendship, masking any potential influence of homophily in affiliation on cooperation. However, in contrast to some previous studies (see [43,97]), we did not find friendships to be driven by personality homophily. This could be attributed to the use of only one proxy, contact sit, and not grooming to define friendship. While combining the two cannot be achieved in this study due to data dependence, future studies should be designed to collect independent measures of these variables to understand the evolutionary origins of homophily better.

In contrast to personality homophily, we found personality heterophily as a proximate mechanism fostering cooperation. Less explorative individuals were likely to form cooperative dyads with more explorative ones. In addition, individuals who differed in the activity-sociability trait were highly successful in cooperation. These findings are in line with our prediction that heterophily in non-social personality traits can positively influence cooperation, potentially due to varying problem-solving perspectives of the associated individuals. From a relatively simple cognitive point of view, dyads with non-social personality differences may represent a ‘leader-follower’ relationship ([98], also see [99]). To illustrate, partners with greater explorative tendencies or higher activity levels may function as ‘catalysts’ (c.f. [54]), while their counterparts with lower explorative behaviour or reduced activities may follow them. This, in contrast to dyads with either two ‘leaders’ or two ‘followers’, may have problems coordinating in a cooperation set-up as the one tested in this study. From an evolutionary viewpoint, such a relationship can enable leaders to gain by imposing their preferences on followers but simultaneously reducing conflicting preferences and the ‘cost of consensus’ [99].

## 5. Conclusion

In summary, we illustrated that friendships can strongly predict cooperative dyad formation and its success. Lower disparity in dominance ranks can lead to higher cooperative task success. We also elucidated the potential reasons behind the limited predictive power of kinship and social tolerance concerning cooperation. Finally, we tested the hypothesis that personality homophily and heterophily may act independently at the trait level to drive cooperation. Yet, we only found support for heterophily in the non-social traits linked to cooperation. To conclude, we provided a comprehensive account of some key proximate mechanisms of cooperation and its decision rules (for partner choice) in a group-living primate species. We specifically emphasised the low cognitive demands of these mechanisms yet highlighted their broad evolutionary implications for a better understanding of the evolution of cooperation.

## Supporting information

Electronic supplementary material

## Ethics

The BPRC is accredited by the Association for Assessment and Accreditation of Laboratory Animal Care (AAALAC) and licensed to keep non-human primates for ethological and biomedical research. The institution follows high standards of animal welfare and refinement measures. Our study complied with all ethical regulations and guidelines for animal testing of BPRC’s Animal Experiments Committee and Animal Welfare Organisation (Animal Welfare Organisation/ IvD approval no. 019C and 019E). All the experimenters were trained and certified (FELASA Accredited Laboratory Animal Science 066/19AF) to conduct animal experiments in accordance with the requirements of Article 9 of the Dutch Experiments on Animals Act.

## Data accessibility

Data and codes generated from the study will be made open upon publication.

## Authors’ contributions

D.B.: conceptualisation, data curation, formal analysis, investigation, methodology, resources, visualisation, writing—original draft, writing—review and editing; S.W.: investigation, methodology, writing—review and editing; E.M.: data curation; E.H.M.S.: methodology, project administration, resources, writing—review and editing.; J.J.M.M.: conceptualisation, methodology, project administration, resources, supervision, writing— review and editing. All authors gave their final approval for publication and agreed to be held accountable for the work performed therein.

## Conflict of interest declaration

We have no competing interests.

## Funding

The study received funding from the European Union’s Horizon 2020 Marie Skłodowska-Curie Actions research and innovation program under grant number H2020-MSCA-IF-2019-893016 (awarded to D.B.).

## Acknowledgements

We thank all caretakers at the BPRC for their help and assistance during the study.

## Notes

### Competing Interest Statement

The authors have declared no competing interest.

## References

1. Brosnan SF, de Waal FBM. 2002 A proximate perspective on reciprocal altruism. Human Nature 13, 129–152. (doi:10.1007/s12110-002-1017-2)

2. Packer C, Ruttan L. 1988 The Evolution of Cooperative Hunting. Am Nat 132, 159–198. (doi:10.1086/284844)

3. Clutton-Brock T. 2002 Breeding Together: Kin Selection and Mutualism in Cooperative Vertebrates. Science (1979) 296, 69–72. (doi:10.1126/science.296.5565.69)

4. Burkart JM et al. 2014 The evolutionary origin of human hyper-cooperation. Nat Commun 5, 4747. (doi:10.1038/ncomms5747)

5. Dugatkin LA. 2002 Cooperation in animals: An evolutionary overview. Biol Philos 17, 459–476. (doi:10.1023/A:1020573415343)

6. Bshary R, Oliveira RF. 2015 Cooperation in animals: toward a game theory within the framework of social competence. Curr Opin Behav Sci 3, 31–37. (doi:10.1016/j.cobeha.2015.01.008)

7. Massen JJM, Behrens F, Martin JS, Stocker M, Brosnan SF. 2019 A comparative approach to affect and cooperation. Neurosci Biobehav Rev 107, 370–387. (doi:10.1016/j.neubiorev.2019.09.027)

8. Nowak MA. 2006 Five Rules for the Evolution of Cooperation. Science (1979) 314, 1560–1563. (doi:10.1126/science.1133755)

9. Raihani NJ, Thornton A, Bshary R. 2012 Punishment and cooperation in nature. Trends Ecol Evol 27, 288–295. (doi:10.1016/j.tree.2011.12.004)

10. Egas M, Riedl A. 2008 The economics of altruistic punishment and the maintenance of cooperation. Proceedings of the Royal Society B: Biological Sciences 275, 871–878. (doi:10.1098/rspb.2007.1558)

11. Hamilton WD. 1964 The genetical evolution of social behaviour. II. J Theor Biol 7, 17–52. (doi:10.1016/0022-5193(64)90039-6)

12. Trivers RL. 1971 The Evolution of Reciprocal Altruism. Q Rev Biol 46, 35–57. (doi:10.1086/406755)

13. Wedekind C, Milinski M. 1996 Human cooperation in the simultaneous and the alternating Prisoner’s Dilemma: Pavlov versus Generous Tit-for-Tat. Proceedings of the National Academy of Sciences 93, 2686–2689. (doi:10.1073/pnas.93.7.2686)

14. Nowak M, Sigmund K. 1993 A strategy of win-stay, lose-shift that outperforms tit-for-tat in the Prisoner’s Dilemma game. Nature 364, 56–58. (doi:10.1038/364056a0)

15. de Waal FBM, Luttrell LM. 1988 Mechanisms of social reciprocity in three primate species: Symmetrical relationship characteristics or cognition? Ethol Sociobiol 9, 101–118. (doi:10.1016/0162-3095(88)90016-7)

16. Stevens JR, Hauser MD. 2004 Why be nice? Psychological constraints on the evolution of cooperation. Trends Cogn Sci 8, 60–65. (doi:10.1016/j.tics.2003.12.003)

17. de Waal FBM. 2000 Attitudinal reciprocity in food sharing among brown capuchin monkeys. Anim Behav 60, 253–261. (doi:10.1006/anbe.2000.1471)

18. Schino G, Aureli F. 2009 Chapter 2 Reciprocal Altruism in Primates. pp. 45–69. (doi:10.1016/S0065-3454(09)39002-6)

19. Zijlstra TW, de Vries H, Sterck EHM. 2021 Emotional bookkeeping and differentiated affiliative relationships: Exploring the role of dynamics and speed in updating relationship quality in the EMO-model. PLoS One 16, e0249519. (doi:10.1371/journal.pone.0249519)

20. Evers E, de Vries H, Spruijt BM, Sterck EHM. 2014 The EMO-Model: An Agent-Based Model of Primate Social Behavior Regulated by Two Emotional Dimensions, Anxiety-FEAR and Satisfaction-LIKE. PLoS One 9, e87955. (doi:10.1371/journal.pone.0087955)

21. Evers E, de Vries H, Spruijt BM, Sterck EHM. 2015 Emotional Bookkeeping and High Partner Selectivity Are Necessary for the Emergence of Partner-Specific Reciprocal Affiliation in an Agent-Based Model of Primate Groups. PLoS One 10, e0118921. (doi:10.1371/journal.pone.0118921)

22. Evers E, de Vries H, Spruijt BM, Sterck EHM. 2016 Intermediate-term emotional bookkeeping is necessary for long-term reciprocal grooming partner preferences in an agent-based model of macaque groups. PeerJ 4, e1488. (doi:10.7717/peerj.1488)

23. Campennì M, Schino G. 2016 Symmetry-based reciprocity: evolutionary constraints on a proximate mechanism. PeerJ 4, e1812. (doi:10.7717/peerj.1812)

24. Puga-Gonzalez I, Ostner J, Schülke O, Sosa S, Thierry B, Sueur C. 2018 Mechanisms of reciprocity and diversity in social networks: a modeling and comparative approach. Behavioral Ecology 29, 745–760. (doi:10.1093/beheco/ary034)

25. De Waal FBM, Luttrell LM. 1989 Toward a comparative socioecology of the genusMacaca: Different dominance styles in rhesus and stumptail monkeys. Am J Primatol 19, 83–109. (doi:10.1002/ajp.1350190203)

26. Hare B, Wobber V, Wrangham R. 2012 The self-domestication hypothesis: evolution of bonobo psychology is due to selection against aggression. Anim Behav 83, 573–585. (doi:10.1016/j.anbehav.2011.12.007)

27. Hare B, Melis AP, Woods V, Hastings S, Wrangham R. 2007 Tolerance Allows Bonobos to Outperform Chimpanzees on a Cooperative Task. Current Biology 17, 619–623. (doi:10.1016/j.cub.2007.02.040)

28. Asakawa-Haas K, Schiestl M, Bugnyar T, Massen JJM. 2016 Partner Choice in Raven (Corvus corax) Cooperation. PLoS One 11, e0156962. (doi:10.1371/journal.pone.0156962)

29. Fruteau C, Voelkl B, van Damme E, Noë R. 2009 Supply and demand determine the market value of food providers in wild vervet monkeys. Proceedings of the National Academy of Sciences 106, 12007–12012. (doi:10.1073/pnas.0812280106)

30. Majolo B, Ames K, Brumpton R, Garratt R, Hall K, Wilson N. 2006 Human friendship favours cooperation in the Iterated Prisoner’s Dilemma. Behaviour 143, 1383–1395. (doi:10.1163/156853906778987506)

31. Silk JB, Alberts SC, Altmann J, Cheney DL, Seyfarth RM. 2012 Stability of partner choice among female baboons. Anim Behav 83, 1511–1518. (doi:10.1016/j.anbehav.2012.03.028)

32. De Moor D, Roos C, Ostner J, Schülke O. 2020 Bonds of bros and brothers: Kinship and social bonding in postdispersal male macaques. Mol Ecol 29, 3346–3360. (doi:10.1111/mec.15560)

33. Silk J, Cheney D, Seyfarth R. 2013 A practical guide to the study of social relationships. Evolutionary Anthropology: Issues, News, and Reviews 22, 213–225. (doi:10.1002/evan.21367)

34. Brent LJN, Chang SWC, Gariépy J-F, Platt ML. 2014 The neuroethology of friendship. Ann N Y Acad Sci 1316, 1–17. (doi:10.1111/nyas.12315)

35. Barclay P. 2016 Biological markets and the effects of partner choice on cooperation and friendship. Curr Opin Psychol 7, 33–38. (doi:10.1016/j.copsyc.2015.07.012)

36. Jehn KA, Shah PP. 1997 Interpersonal relationships and task performance: An examination of mediation processes in friendship and acquaintance groups. J Pers Soc Psychol 72, 775–790. (doi:10.1037/0022-3514.72.4.775)

37. Réale D, Reader SM, Sol D, McDougall PT, Dingemanse NJ. 2007 Integrating animal temperament within ecology and evolution. Biological Reviews 82, 291–318. (doi:10.1111/j.1469-185X.2007.00010.x)

38. Moiron M, Laskowski KL, Niemelä PT. 2020 Individual differences in behaviour explain variation in survival: a meta-analysis. Ecol Lett 23, 399–408. (doi:10.1111/ele.13438)

39. Smith BR, Blumstein DT. 2008 Fitness consequences of personality: a meta-analysis. Behavioral Ecology 19, 448–455. (doi:10.1093/beheco/arm144)

40. Verspeek J, Staes N, Leeuwen EJC van, Eens M, Stevens JMG. 2019 Bonobo personality predicts friendship. Sci Rep 9, 19245. (doi:10.1038/s41598-019-55884-3)

41. Laakasuo M, Rotkirch A, van Duijn M, Berg V, Jokela M, David-Barrett T, Miettinen A, Pearce E, Dunbar R. 2020 Homophily in Personality Enhances Group Success Among Real-Life Friends. Front Psychol 11. (doi:10.3389/fpsyg.2020.00710)

42. Noë N, Whitaker RM, Chorley MJ, Pollet T V. 2016 Birds of a feather locate together? Foursquare checkins and personality homophily. Comput Human Behav 58, 343–353. (doi:10.1016/j.chb.2016.01.009)

43. Massen JJM, Koski SE. 2014 Chimps of a feather sit together: chimpanzee friendships are based on homophily in personality. Evolution and Human Behavior 35, 1–8. (doi:10.1016/j.evolhumbehav.2013.08.008)

44. Graziano WG, Eisenberg N. 1997 Agreeableness. In Handbook of Personality Psychology, pp. 795–824. Elsevier. (doi:10.1016/B978-012134645-4/50031-7)

45. Lukaszewski AW, von Rueden CR. 2015 The extraversion continuum in evolutionary perspective: A review of recent theory and evidence. Pers Individ Dif 77, 186–192. (doi:10.1016/j.paid.2015.01.005)

46. Molesti S, Majolo B. 2016 Cooperation in wild Barbary macaques: factors affecting free partner choice. Anim Cogn 19, 133–146. (doi:10.1007/s10071-015-0919-4)

47. Koski SE, Burkart JM. 2015 Common marmosets show social plasticity and group-level similarity in personality. Sci Rep 5, 8878. (doi:10.1038/srep08878)

48. Schuett W, Dall SRX, Royle NJ. 2011 Pairs of zebra finches with similar ‘personalities’ make better parents. Anim Behav 81, 609–618. (doi:10.1016/j.anbehav.2010.12.006)

49. Spoon TR, Millam JR, Owings DH. 2006 The importance of mate behavioural compatibility in parenting and reproductive success by cockatiels, Nymphicus hollandicus. Anim Behav 71, 315–326. (doi:10.1016/j.anbehav.2005.03.034)

50. LePine JA, Van Dyne L. 2001 Voice and cooperative behavior as contrasting forms of contextual performance: Evidence of differential relationships with Big Five personality characteristics and cognitive ability. Journal of Applied Psychology 86, 326–336. (doi:10.1037/0021-9010.86.2.326)

51. Reilly RR, Lynn GS, Aronson ZH. 2002 The role of personality in new product development team performance. Journal of Engineering and Technology Management 19, 39–58. (doi:10.1016/S0923-4748(01)00045-5)

52. Mohammed S, Angell LC. 2003 Personality Heterogeneity in Teams. Small Group Res 34, 651–677. (doi:10.1177/1046496403257228)

53. Peeters MAG, van Tuijl HFJM, Rutte CG, Reymen IMMJ. 2006 Personality and team performance: a meta-analysis. Eur J Pers 20, 377–396. (doi:10.1002/per.588)

54. Scheid C, Noë R. 2010 The performance of rooks in a cooperative task depends on their temperament. Anim Cogn 13, 545–553. (doi:10.1007/s10071-009-0305-1)

55. Gilby IC, Eberly LE, Wrangham RW. 2008 Economic profitability of social predation among wild chimpanzees: individual variation promotes cooperation. Anim Behav 75, 351–360. (doi:10.1016/j.anbehav.2007.06.008)

56. Scherer U, Kuhnhardt M, Schuett W. 2017 Different or alike? Female rainbow kribs choose males of similar consistency and dissimilar level of boldness. Anim Behav 128, 117–124. (doi:10.1016/j.anbehav.2017.04.007)

57. Dingemanse NJ, Both C, Drent PJ, Tinbergen JM. 2004 Fitness consequences of avian personalities in a fluctuating environment. Proc R Soc Lond B Biol Sci 271, 847–852. (doi:10.1098/rspb.2004.2680)

58. Koski SE. 2011 Social personality traits in chimpanzees: temporal stability and structure of behaviourally assessed personality traits in three captive populations. Behav Ecol Sociobiol 65, 2161–2174. (doi:10.1007/s00265-011-1224-0)

59. Thierry B. 2022 Where do we stand with the covariation framework in primate societies? American Journal of Biological Anthropology 178, 5–25. (doi:10.1002/ajpa.24441)

60. Massen JJM, van den Berg LM, Spruijt BM, Sterck EHM. 2010 Generous Leaders and Selfish Underdogs: Pro-Sociality in Despotic Macaques. PLoS One 5, e9734. (doi:10.1371/journal.pone.0009734)

61. Massen JJM, Luyten IJAF, Spruijt BM, Sterck EHM. 2011 Benefiting friends or dominants: prosocial choices mainly depend on rank position in long-tailed macaques (Macaca fascicularis). Primates 52, 237–247. (doi:10.1007/s10329-011-0244-8)

62. van Leeuwen EJC, DeTroy SE, Kaufhold SP, Dubois C, Schütte S, Call J, Haun DBM. 2021 Chimpanzees behave prosocially in a group-specific manner. Sci Adv 7. (doi:10.1126/sciadv.abc7982)

63. Sigmundson R, Stribos MS, Hammer R, Herzele J, Pflüger LS, Massen JJM. 2021 Exploring the Cognitive Capacities of Japanese Macaques in a Cooperation Game. Animals 11, 1497. (doi:10.3390/ani11061497)

64. Sterck EHM, Olesen CU, Massen JJM. 2015 No costly prosociality among related long-tailed macaques (Macaca fascicularis). J Comp Psychol 129, 275–282. (doi:10.1037/a0039180)

65. Stocker M, Loretto M-C, Sterck EHM, Bugnyar T, Massen JJM. 2020 Cooperation with closely bonded individuals reduces cortisol levels in long-tailed macaques. R Soc Open Sci 7, 191056. (doi:10.1098/rsos.191056)

66. Bhattacharjee D, Cousin E, Pflüger LS, Massen JJM. 2023 Prosociality in a despotic society. iScience 26, 106587. (doi:10.1016/j.isci.2023.106587)

67. Phillips T. 2018 The concepts of asymmetric and symmetric power can help resolve the puzzle of altruistic and cooperative behaviour. Biological Reviews 93, 457–468. (doi:10.1111/brv.12352)

68. Hirata S, Fuwa K. 2007 Chimpanzees (Pan troglodytes) learn to act with other individuals in a cooperative task. Primates 48, 13–21. (doi:10.1007/s10329-006-0022-1)

69. Massen JJM, Ritter C, Bugnyar T. 2015 Tolerance and reward equity predict cooperation in ravens (Corvus corax). Sci Rep 5, 15021. (doi:10.1038/srep15021)

70. Martin JS, Koski SE, Bugnyar T, Jaeggi AV, Massen JJM. 2021 Prosociality, social tolerance and partner choice facilitate mutually beneficial cooperation in common marmosets, Callithrix jacchus. Anim Behav 173, 115–136. (doi:10.1016/j.anbehav.2020.12.016)

71. Martin JS, Koski SE, Bugnyar T, Jaeggi AV, Massen JJM. 2021 Prosociality, social tolerance and partner choice facilitate mutually beneficial cooperation in common marmosets, Callithrix jacchus. Anim Behav 173, 115–136. (doi:10.1016/j.anbehav.2020.12.016)

72. Bhattacharjee D, Rut Guðjónsdóttir A, Escriche Chova P, Jäckels J, de Groot NG, Wallner B, Massen JJ, Pflüger LS. 2023 Behavioural, physiological, and genetic drivers of coping. bioRxiv (doi:10.1101/2023.08.28.555090)

73. Kluiver CE, de Jong JA, Massen JJM, Bhattacharjee D. 2022 Personality as a Predictor of Time-Activity Budget in Lion-Tailed Macaques (Macaca silenus). Animals 12, 1495. (doi:10.3390/ani12121495)

74. Neumann C, Fischer J. 2023 Extending Bayesian Elo-rating to quantify the steepness of dominance hierarchies. Methods Ecol Evol 14, 669–682. (doi:10.1111/2041-210X.14021)

75. Schülke O et al. 2022 Quantifying within-group variation in sociality—covariation among metrics and patterns across primate groups and species. Behav Ecol Sociobiol 76, 50. (doi:10.1007/s00265-022-03133-5)

76. R Development Core Team. 2019 R Core Team (2020). R: A language and environment for statistical computing. R Foundation for Statistical Computing, Vienna, Austria. URL https://www.R-project.org/. R Foundation for Statistical Computing. 2.

77. Brooks ME, Kristensen K, Benthem KJ, van Magnusson A, Berg CW, Nielsen A, Skaug HJ, Mächler M, Bolker BM. 2017 glmmTMB Balances Speed and Flexibility Among Packages for Zero-inflated Generalized Linear Mixed Modeling. R J 9, 378. (doi:10.32614/RJ-2017-066)

78. Lüdecke D, Ben-Shachar M, Patil I, Waggoner P, Makowski D. 2021 performance: An R Package for Assessment, Comparison and Testing of Statistical Models. J Open Source Softw 6, 3139. (doi:10.21105/joss.03139)

79. Hothorn T, Zeileis A. 2011 Diagnostic Checking in Regression Relationships. R News 2.

80. Hartig F. 2020 DHARMa: Residual Diagnostics for Hierarchical Regression Models. The Comprehensive R Archive Network.

81. van Leeuwen EJC, DeTroy SE, Kaufhold SP, Dubois C, Schütte S, Call J, Haun DBM. 2021 Chimpanzees behave prosocially in a group-specific manner. Sci Adv 7. (doi:10.1126/sciadv.abc7982)

82. Ruiter JRD, Geffen E. 1998 Relatedness of matrilines, dispersing males and social groups in long–tailed macaques (Macaca fascicularis). Proc R Soc Lond B Biol Sci 265, 79–87. (doi:10.1098/rspb.1998.0267)

83. Sterck EHM, Watts DP, van Schaik CP. 1997 The evolution of female social relationships in nonhuman primates. Behav Ecol Sociobiol 41, 291–309. (doi:10.1007/s002650050390)

84. Schülke O, Bhagavatula J, Vigilant L, Ostner J. 2010 Social Bonds Enhance Reproductive Success in Male Macaques. Current Biology 20, 2207–2210. (doi:10.1016/j.cub.2010.10.058)

85. Silk JB. 2009 Nepotistic cooperation in non-human primate groups. Philosophical Transactions of the Royal Society B: Biological Sciences 364, 3243–3254. (doi:10.1098/rstb.2009.0118)

86. Chapais B, Prud’Homme J, Teijeiro S. 1994 Dominance competition among siblings in Japanese macaques: constraints on nepotism. Anim Behav 48, 1335–1347. (doi:10.1006/anbe.1994.1370)

87. Langergraber KE, Mitani JC, Vigilant L. 2007 The limited impact of kinship on cooperation in wild chimpanzees. Proceedings of the National Academy of Sciences 104, 7786–7790. (doi:10.1073/pnas.0611449104)

88. Schaub H. 1996 Testing kin altruism in long-tailed macaques (Macaca fascicularis) in a food-sharing experiment. Int J Primatol 17, 445–467. (doi:10.1007/BF02736631)

89. Suchak M, Eppley TM, Campbell MW, de Waal FBM. 2014 Ape duos and trios: spontaneous cooperation with free partner choice in chimpanzees. PeerJ 2, e417. (doi:10.7717/peerj.417)

90. Schwing R, Meaux E, Piseddu A, Huber L, Noë R. 2021 Kea, Nestor notabilis, achieve cooperation in dyads, triads, and tetrads when dominants show restraint. Learn Behav 49, 36–53. (doi:10.3758/s13420-021-00462-9)

91. Sinha A. 1998 Knowledge acquired and decisions made: triadic interactions during allogrooming in wild bonnet macaques, Macaca radiata. Philos Trans R Soc Lond B Biol Sci 353, 619–631. (doi:10.1098/rstb.1998.0230)

92. Sueur C, Petit O, De Marco A, Jacobs AT, Watanabe K, Thierry B. 2011 A comparative network analysis of social style in macaques. Anim Behav 82, 845–852. (doi:10.1016/j.anbehav.2011.07.020)

93. Seyfarth RM, Cheney DL. 2012 The Evolutionary Origins of Friendship. Annu Rev Psychol 63, 153–177. (doi:10.1146/annurev-psych-120710-100337)

94. Massen J, Sterck E, de Vos H. 2010 Close social associations in animals and humans: functions and mechanisms of friendship. Behaviour 147, 1379–1412. (doi:10.1163/000579510X528224)

95. Stocker M, Loretto M-C, Sterck EHM, Bugnyar T, Massen JJM. 2020 Cooperation with closely bonded individuals reduces cortisol levels in long-tailed macaques. R Soc Open Sci 7, 191056. (doi:10.1098/rsos.191056)

96. Leeuwen EJC, Van Donink S, Eens M, Stevens JMG. 2021 Group-level variation in co-feeding tolerance between two sanctuary-housed communities of chimpanzees (Pan troglodytes). Ethology 127, 517–526. (doi:10.1111/eth.13154)

97. Ebenau A, von Borell C, Penke L, Ostner J, Schülke O. 2019 Personality homophily affects male social bonding in wild Assamese macaques, Macaca assamensis. Anim Behav 155, 21–35. (doi:10.1016/j.anbehav.2019.05.020)

98. Johnstone RA, Manica A. 2011 Evolution of personality differences in leadership. Proceedings of the National Academy of Sciences 108, 8373–8378. (doi:10.1073/pnas.1102191108)

99. Conradt L, Roper TJ. 2009 Conflicts of interest and the evolution of decision sharing. Philosophical Transactions of the Royal Society B: Biological Sciences 364, 807–819. (doi:10.1098/rstb.2008.0257)

