## Supplementary material for "Personality heterophily and friendship as drivers for successful cooperation": Electronic supplementary material

*Electronic Supplementary Material to*

**Table of contents**

| Item | Description | Page |
| --- | --- | --- |
| Table S1 | Details of the individuals in the study | 02 |
| Movie S1 | A cooperation trial from the testing phase | 02 |

**Table S1**

| **Identity and sex of the individuals** | | | **Date of birth** |
| --- | --- | --- | --- |
| **Group 1** | | | |
| Dukki ♀ |  |  | 13/10/2017 |
| Mooi ♀ |  |  | 16/08/2017 |
| Alibi ♀ |  |  | 18/10/2021 |
| Toffi ♀ |  |  | 30/09/2017 |
| Walibi ♀ |  |  | 10/09/2012 |
|  | Bowi ♀ |  | 18/10/2020 |
| Urbi-et-orbi ♀ |  |  | 13/10/2014 |
|  | Tutudetuu ♂ |  | 11/11/2018 |
|  | Oui ♀ |  | 18/09/2020 |
| Tebbi ♀ |  |  | 31/08/2017 |
|  | Impromptu ♂ |  | 29/01/2021 |
| **Group 2** | | | |
| Rizzo ♂ |  |  | 24/10/2009 |
| Castello ♀ |  |  | 14/01/2008 |
| Saboteur ♀ |  |  | 17/08/2007 |
|  | Tamayo ♀ |  | 20/10/2017 |
| Hacienda ♀ |  |  | 19/01/2008 |
| Magelaen ♀ |  |  | 14/08/2007 |
|  | Soest ♀ |  | 11/05/2015 |
|  |  | Emerald ♀ | 09/04/2019 |
|  |  | Freggel ♂ | 09/04/2020 |
|  |  | Elmo ♂ | 15/02/2021 |
|  | Etten ♀ |  | 13/08/2016 |
|  | Rox ♀ |  | 18/04/2017 |
|  | Lageveen ♀ |  | 17/10/2018 |
|  | Monopoly ♂ |  | 08/06/2019 |
|  | Stas ♀ |  | 21/08/2014 |
|  |  | Nellie ♀ | 14/06/2019 |
|  |  | Sjors ♂ | 27/02/2021 |
| **Group 3** | | | |
| Sollie ♂ |  |  | 12/12/2015 |
| Horsten ♂ |  |  | 20/10/2017 |
| Kluivers ♂ |  |  | 22/10/2017 |
| Driel ♂ |  |  | 04/12/2017 |

**Table S1 – Details of the individuals in the study.** The identity, sex, relatedness, and date of birth of the participating individuals from the three groups are summarised. The placement of the macaques from left to right indicates mother-offspring relationships. For example, Bowi is the daughter of Walibi in Gr.1.

**Movie S1 – A cooperation trial from the testing phase.** A trial from the testing phase showing the cooperative success of a long-tailed macaque dyad.
